# Sex-specific niche signaling contributes to sexual dimorphism following stem cell transplantation

**DOI:** 10.1101/2022.06.13.495897

**Authors:** Julianne N.P. Smith, Brittany A. Cordova, Brian Richardson, Kelsey F. Christo, Jordan Campanelli, Alyssia V. Broncano, Jonathan Chen, Juyeun Lee, Scott J. Cameron, Justin D. Lathia, Wendy A. Goodman, Mark J. Cameron, Amar B. Desai

**Author notes:** Correspondence: Amar B. Desai, Department of Medicine and Case Comprehensive Cancer Center, Case Western Reserve University.

## Abstract

Hematopoietic stem cell (HSC) transplantation (HST) is a curative treatment for many hematopoietic cancers and bone marrow (BM) disorders but is currently limited by numerous complications including a lengthy recovery period, prolonged neutropenia resulting in severe infections and bleeding, and a high incidence of graft vs. host disease (GVHD). While clinical studies have demonstrated that sex mismatch, notably male recipients with female donor cells, results in increased risk of GVHD (likely due to male recipient minor histocompatibility antigens targeted by donor female T-cells ^1^), increased non-relapse mortality, and decreased overall survival, the mechanisms underlying sex-determinants on hematopoiesis and post-transplant recovery are not clear. In this manuscript we have identified: 1) unique expression of hematopoietic niche factors in the BM and spleens of male and female mice, 2) altered kinetics of hematopoietic reconstitution following transplantation when male vs. female BM is used as the donor cell source, 3) a sex-specific role for the recipient niche in promoting post HST recovery, and 4) a dose-dependent role for exogenous sex hormones in maintaining hematopoietic stem and progenitor cells (HSPCs). Taken together, these data demonstrate that sex-specific cellular and molecular signaling occurs during hematopoietic regeneration. Further identifying novel sex-dependent determinants of regeneration following transplantation will not only enhance understanding of steady state versus regeneration hematopoiesis but may also reveal unique (and potentially sex-specific) therapeutic targets to accelerate hematologic recovery.

**Key Points:**

1. Male and female mice display altered kinetics of regeneration following HST due to unique niche factors in hematopoietic compartments.
2. Exogenous steroid sex hormones uniquely regulate the pool of hematopoietic stem and progenitor cells and may impact transplantation outcomes.

## Introduction

Hematopoietic stem cells (HSCs) are derived from the bone marrow (BM) and tasked with ensuring a consistent output of differentiated blood cell types throughout the lifetime of an organism. This involves balancing the acts of self-renewal, differentiation to mature progeny, and proliferation in response to infection, stress, or injury to provide lifelong hematopoietic function. At homeostasis, HSCs maintain genomic stability and avoid replicative stress by existing in a quiescent state, enforced by both HSC intrinsic and microenvironmental cues. During stress, which can be triggered by infection, inflammation, bleeding, certain medications, exposure to irradiation or chemotherapy, or by procedures such as hematopoietic stem cell transplantation (HST), a regenerative hematopoiesis program occurs in which HSCs receive specific signals to activate and produce progenitor populations capable of restoring homeostasis through mobilization and differentiation to replenish depleted populations. Elucidating the specific signaling pathways associated with regenerative hematopoiesis has implications for the approximately ~25,000 HST are performed each year in the United States to treat conditions including several plasma cell dyscrasias – most of which constitute multiple myeloma, non-Hodgkin’s lymphoma, acute myeloid leukemia, myelodysplastic syndrome, and acute lymphocytic leukemia.

Current efforts to improve HST efficacy involve minimizing both recipient risk factors (including diagnosis stage, time to transplant, age, and opportunistic infections) and donor risk factors (including HLA mismatch, age, KIR genotype, and sex mismatch)^2^. While recent literature has identified intrinsic differences in hematopoietic properties between males and females, including observing reduced frequencies of circulating hematopoietic progenitor cells in females versus males^3^ and elucidating critical roles for gonad derived sex hormones in maintenance of HSC self-renewal and proliferative capacity,^4 5^ major gaps exist in understanding sexual dimorphism in the context of hematopoietic regeneration. Additionally, with our lab’s recent work implicating a role for the splenic niche in HST biology,^6^ defining the sex-dependent role of each hematopoietic compartment in post-transplant hematopoiesis remains a priority for not only fundamental understanding of HST, but also its clinical use. Here we use a variety of *in vitro* culture models along with *in vivo* transplantation assays to provide novel biological insights into sex-specific drivers of regeneration, which has significant potential to uncover sex-specific therapeutic targets to promote HST recovery.

## Methods

### Animals

Animals were housed in the AAALAC accredited facilities of the CWRU School of Medicine. Husbandry and experimental procedures were approved by the Case Western Reserve University Institutional Animal Care and Use Committee (IACUC) in accordance with approved IACUC protocol 2019-0065. Steady-state and transplantation analyses were performed on 8wk old female C57BL/6J mice obtained from Jackson Laboratories. B6.SJL-Ptprca Pepcb/BoyJ and splenectomized C57BL/6 mice were obtained from Jackson Laboratories. All animals were observed daily for signs of illness. Mice were housed in standard microisolator cages and maintained on a defined, irradiated diet and autoclaved water.

### Complete Blood Count Analysis

Peripheral blood was collected into Microtainer EDTA tubes (Becton-Dickinson) by submandibular cheek puncture. Blood counts were analyzed using a Hemavet 950 FS hematology analyzer.

### Colony Forming Analysis

5e4 total bone marrow cells and 2.5e5 total splenocytes were plated in Methocult media M3434 (StemCell Technologies) supplemented with hemin. CFU-GM and BFU-E colonies were scored 12 days post-plating.

### RNA Extraction, qPCR, and Bulk RNA Sequencing

Total splenocytes and bone marrow cells were collected, lysed and RNA was extracted using the RNeasy MiniKit (QIAGEN) with on-column DNase treatment, according to the manufacturer’s protocol. cDNA was synthesized using the PrimeScript RT Reagent Kit (Takara) following manufacturer’s instructions. Real time PCR measurement was performed in a 20ul reaction containing 1ul cDNA template and a 1:20 dilution of primer/probe with 1X Accuris Taq DNA polymerase. Samples were run on a CFX96 optical module (Bio-Rad). Thermal cycling conditions were 95C for 3 minutes, followed by 50 cycles of 95C for 15 seconds and 60C for 1 minute. Murine probe/primer sets for all genes assayed were obtained from Life Technologies. For each reverse transcription reaction, Cq values were determined as the average values obtained from three independent real-time PCR reactions. For RNAsequencing studies, samples were shipped on dry ice to MedGenome for subsequent library preparation and PE150 sequencing on Illumina (40M reads/sample). Data was analyzed in the Applied Functional Genomics core facility at CWRU. Microarray data is available at the NCBI GEO.

### Bone Marrow Transplantation

Mice were exposed to 10Gy total body irradiation from a cesium source. 16-18hrs later, mice received 1e6 whole bone marrow cells by retroorbital injection.

### Quantification of HSPCs and Splenic Cell Types

Bone marrow cells were obtained by flushing hindlimb bones and splenocytes were obtained by mincing spleens. Cellularity was measured following red blood cell lysis. Cells were stained with antibodies against CD45R/B220 (RA3-6B2), CD11b (M1/70), CD3e (500A2), Ly-6G and Ly6C (RB6-8C5), TER-119 (TER-119), CD117 (2B8), F4/80 (Cl:A3-1), Sca1 (D7), c-kit (2B8), CD150 (TC15-12F12.1), CD48 (HM48-1), and data was acquired on an LSRII flow cytometer (BD Biosciences). Analysis was performed on FlowJo software (TreeStar).

### Bone Marrow Cultures

Total BM was flushed from male and female mice, and 5M cells were plated in RPMI 1640 media containing 2% FBS (charcoal stripped), 100ng/mL Thpo, 50ng/mL SCF, and 100ng/mL Flt3. E2 and T (Sigma) were supplemented into cultures at 3, 30, and 300 pg/mL and cultures were harvested for HSPC quantification 5 days post plating.

### Statistical Analysis

All values were tabulated graphically with error bars corresponding to standard error of the means. Analysis was performed using GraphPad Prism software. Unpaired two-tailed Student’s t-test was used to compare groups, unless otherwise noted. For peripheral blood recovery kinetic analysis, 2-way ANOVA was used.

## Results

### Male and female C57BL/6 mice exhibit minor homeostatic differences in BM and splenic hematopoiesis

To first identify potential sex-specific differences in the cellular composition of the three major hematopoietic tissues, we characterized the peripheral blood, BM, and splenic compartments of young (8–10-week-old) male and female C57/Bl6 mice at steady state. At homeostasis we observe no significant differences in peripheral blood outside of a moderate but insignificant increase in male neutrophil counts, and no differences between white blood cells, lymphocytes, red blood cells, and platelets. In the bone marrow we observed minor differences in composition as male mice demonstrated a trend towards increased total cellularity (23M cells/hindlimb vs. 20M cells/hindlimb), a significant increase in the total number of lineage (-), Sca-1(+), c-Kit(+) LSK cells (49k/hindlimb vs. 33k/hindlimb), and a notably but statistically insignificant reduction in the number of long-term (LT)-HSC, as marked by LSK/CD150+/CD48(-), (1600/hindlimb vs. 2200/hindlimb). Importantly, these differences were not likely due to the increased size of the BM compartment in male mice, as LSK frequencies were also higher in males (**Supplemental Figure 1**). In the spleen we observed no significant differences in splenic cellularity or LSK populations (**Figure 1A-C**). Additional measures of colony forming capacity within the BM and splenic compartments demonstrate no difference in the output of burst forming-erythroid or granulocyte-macrophage colonies between male and female mice (**Figure 1D**). Together, these data indicate that male and female mice exhibit generally similar hematopoietic activity and peripheral blood cell output at steady-state.

**Figure 1:**
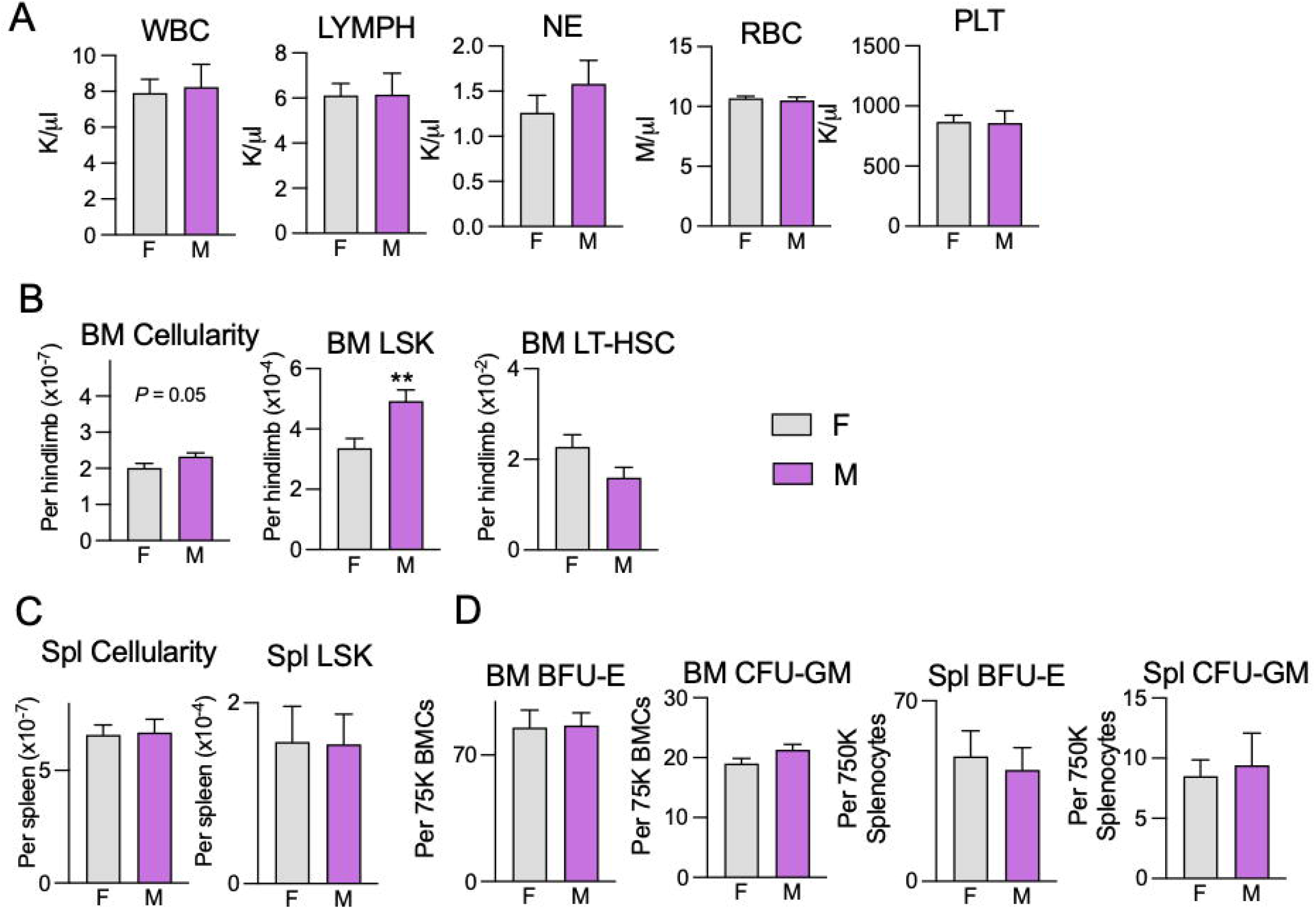
Male and female C57BL/6 mice exhibit minor homeostatic differences in BM and splenic hematopoiesis-. A. Complete blood count analysis of circulating white blood cells (WBC), lymphocytes (LYMPH), neutrophils (NE), red blood cells (RBC), and platelets (PLT) in steady-state female (F) and male (M) mice. B, C. Quantification of total nucleated bone marrow cells flushed per hindlimb and spleen, and immunophenotypic analysis of hematopoietic stem and progenitor cells (HSPCs; Lineage-Sca1+ c-Kit+ (LSK)), and hematopoietic stem cells (HSCs; LSK CD48-CD150+) per hindlimb and per spleen. N=15 mice/arm. D. Number of burst forming unit-erythroid (BFU-E) and colony forming unit-granulocyte-macrophage (CFU-GM) in the marrow and spleen of male and female mice. N=6 mice/arm. Error bars represent SEM. ** *P* < 0.01. Student’s t-test used for all analyses.

### The hematopoietic microenvironments of male and female mice uniquely express hematopoietic niche factors

Given the difference in phenotypic HSPCs we identified at homeostasis, we performed real-time PCR on total BM and splenic RNA from age-matched young male and female animals to evaluate differentially expressed with the hematopoietic niche^7^ genes. Notably in the BM we found expression differences in males as compared to females, including an increase in *Il6* (2.5-fold) coupled with decreases in *Vcam1^8^* (0.36-fold), *Cxcl12^9^* (0.19-fold), and Notch^*10*^ factors (0.17-fold for *Notch1*). The reduction in *Vcam1* and *Cxcl12*, which are associated with HSPC retention in the niche ^11 12 13^, coupled with decreased Notch signaling (which has been implicated in maintenance of HSC quiescence ^14^), suggests the male BM microenvironment is primed for proliferation. Additionally, Il6 has been shown both to enhance the myeloid differentiation of HSPCs^15^, and to promote IL-3 induced hematopoietic proliferation and increased HSC production ^16^, thus also promoting a more active HSC microenvironment. In the spleen we observed additional significant differences including increased male *Pf4^17^* (5.4-fold), *Angpt1^18^* (2.6-fold), and *Notch1* (3.2-fold), coupled with decreased male *KitL^19^(0.46-fold)* and *Vcam1 (0.34-fold)* expression. (**Figure 2**). Thus, contrary to the signature observed in the BM, the significant increases in *Pf4, Angpt1, and Notch1* suggests that the male spleen exists in a more quiescent state compared to female counterparts. The existence of opposing signatures within the splenic and BM microenvironment is intriguing and in line with previous work identifying differential roles for each organ in HSC maintenance. These data, however, advance our understanding of these anatomical differences by uncovering sex-dependent regulators within the two hematopoietic microenvironments.

**Figure 2:**
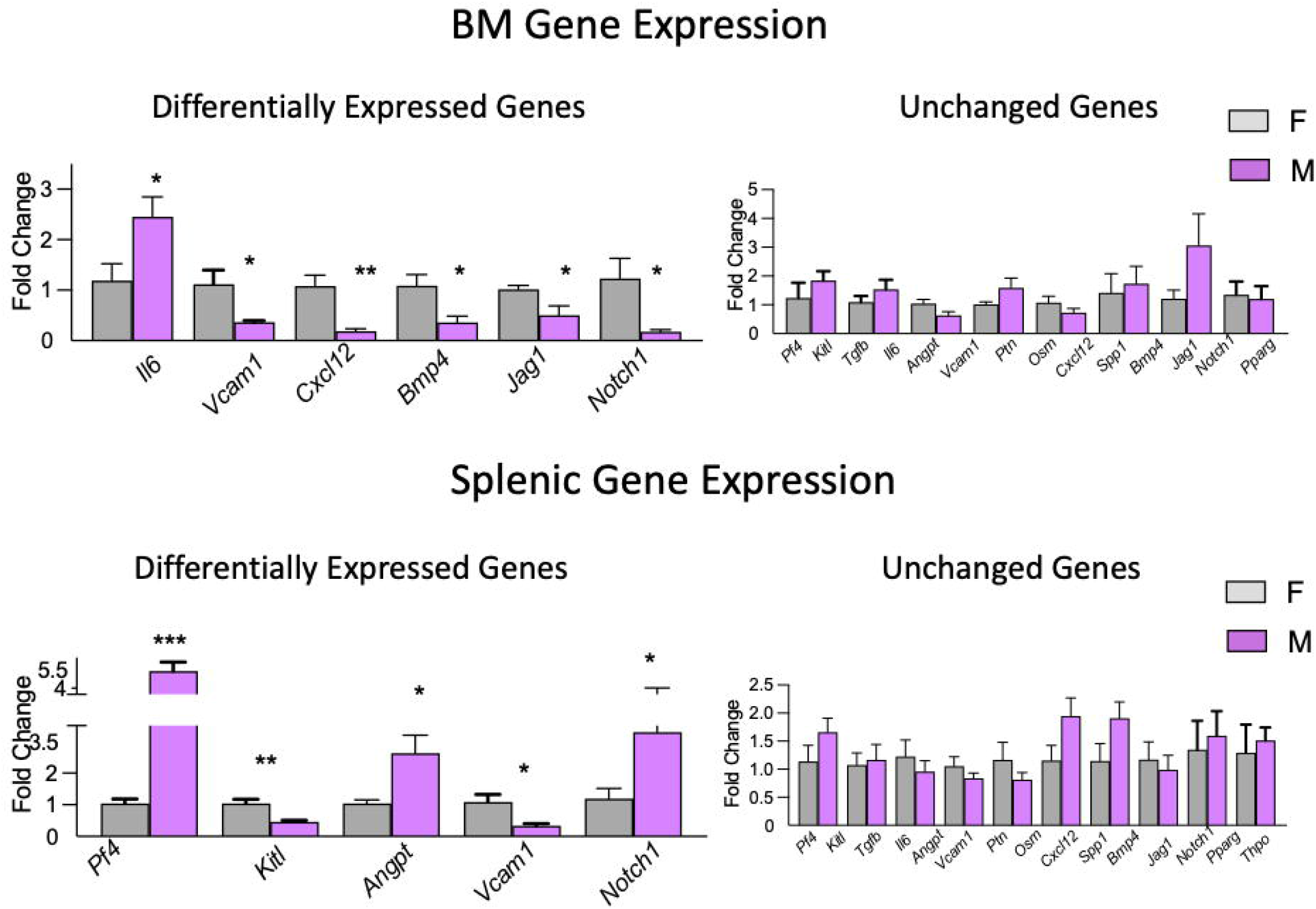
The hematopoietic microenvironments of male and female mice uniquely express hematopoietic niche factors-. Fold-change of indicated genes in total BM and splenic RNA preps from male (purple) over female (gray) mice, normalized to *Actb*. N=5 mice/arm. Error bars represent SEM. \*\**P* < 0.01. Student’s t-test used for all analyses.

### Male and female mice display comparable early rates of peripheral recovery following syngeneic HST

To determine whether sex-dependent gene expression patterns are associated with functional differences, we characterized the kinetics of hematopoietic recovery following HST in syngeneic transplants of young female BM into young female recipients (F➔F) and young male BM into young male recipients (M➔M) (**Figure 3A**). Following lethal irradiation and transplantation of 1e6 total BM cells, we assessed peripheral complete blood counts (CBCs) at Day (D) 7, 14, 21, and 28 post-transplant. We observed only minor changes in white blood cell (WBC) differentials through D21, with the only significant difference being increased female lymphocytes at D14, however we did observe striking increases in total WBC, lymphocytes, and neutrophils in male mice at D28 post HST. We quantified significantly higher red blood cell (RBC) counts in D7 male vs. female mice, however there were subsequently no significant differences at D14, 21 or 28. Finally, we observed transient differences in platelets at D21 and D28, with female mice displaying significantly higher counts at the former and males higher values at the latter timepoint. Taken together this data suggests that peripheral recovery during the first 21 days post HST is comparable between sexes, with recovery curves starting to diverge towards a male advantage by D28 (**Figure 3B**).

**Figure 3:**
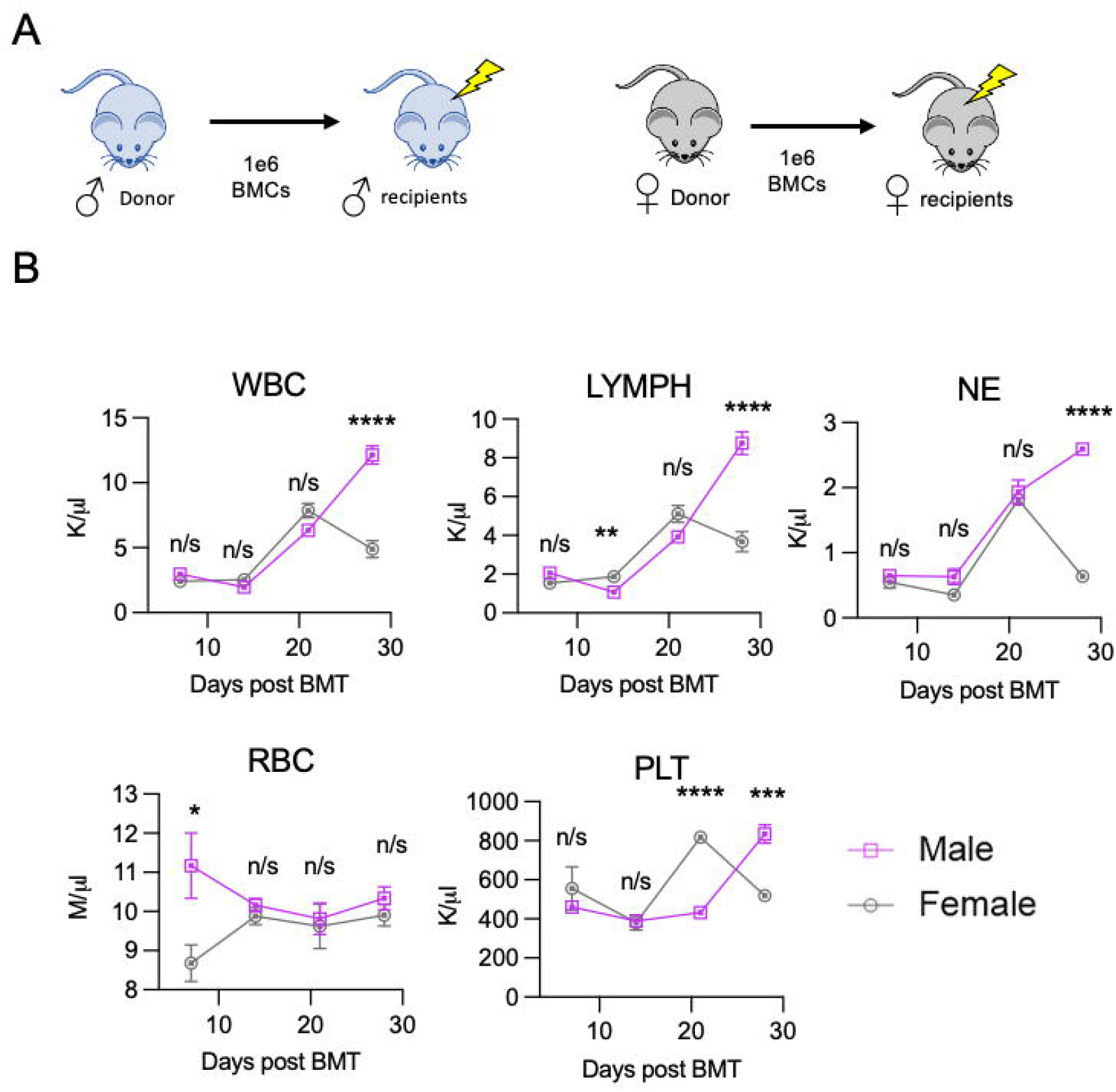
Male and female mice display comparable early rates of peripheral recovery following syngeneic HST-. Schematic depicting transplant conditions. 1e6 BM cells from male donors were transplanted into lethally irradiated (10Gy) male recipients, and the equivalent performed using female donors and recipients. B. Complete blood count analysis of circulating white blood cells (WBC) lymphocytes (LYMPH), neutrophils (NE), red blood cells (RBC), and platelets (PLT) in male (purple) and female (gray) mice at indicated time points post-bone marrow transplantation. N=5 mice/arm for D7, D21 and D28, N=10 mice/arm for D14. Error bars represent SEM. Welch’s T-Test performed at each timepoint.

### Male mice display accelerated medullary and extramedullary hematopoietic recovery following HST

Because the M➔M HST setting demonstrated exuberant myeloid and lymphoid hematopoietic recovery, we next evaluated the kinetics of BM recovery. We observed a significantly accelerated recovery of the M➔M cohort compared to the F➔F mice in terms of total BM cellularity, with cellularity curves beginning to deviate at the D14 (20M cells/hindlimb vs. 17M) timepoint, through D21 (30M cells/hindlimb vs. 20M) and D28 (39M cells/hindlimb vs. 33M) post HST. This was also true for the absolute number of BM LSK cells with the male cohort containing significantly higher numbers of LSK cells at D14 (21K/hindlimb vs. 2K) and D21 (35K/hindlimb vs. 19K), and LT-HSC numbers- with the male cohort containing higher LT-HSC numbers at D14 (650/hindlimb 150), D21 (1750/hindlimb vs. 1250) and D28 (2200 vs. 1050), (**Figure 4B**). Quantification of differentiated cell populations in the BM demonstrated no significant differences in B-cells (as marked by B220+) or T-cells (as marked by CD3e+), however we did observe a significant increase in the number of BM macrophages in male mice (as marked by F480+) at D21 (2.1M vs. 0.83M) and D28 (1.82M vs. 1.24M) (**Figure 4C**). We additionally performed bulk RNA sequencing on total RNA isolated from the BM and spleens of D14 post HST animals (30M paired end reads, MedGenome). Interestingly, in the bone marrow we saw sex differences between many genes associated with hematopoiesis and regeneration, including downregulation of Epas1 whose inhibition has been shown to promote tissue regeneraton^20^, and *Ptn*, which has been shown to promote HSC quiescence following myeloablation^21^), in male compared to female recipients. Differential expression of these genes is consistent with the transient increase in HSPCs we observed in M➔M HST. Additionally, we found concomitant increases in *Ms4a3* (suggesting enhanced regeneration of the granulocyte-monocyte progenitor population ^22^), *Fpr2* (which may promote granulocyte recovery ^23^), *Il1r2* (potentially promoting myeloid cell proliferation ^24^), *Ccr1* and *Hdc* (which can promote myeloid progenitor cell proliferation ^25 26^), and *Ms4a4c* and *Gsr (*potentially promoting alternative activation of macrophages to reduce inflammation in the post HST microenvironment ^27 28^) (**Figure 4D).**

**Figure 4:**
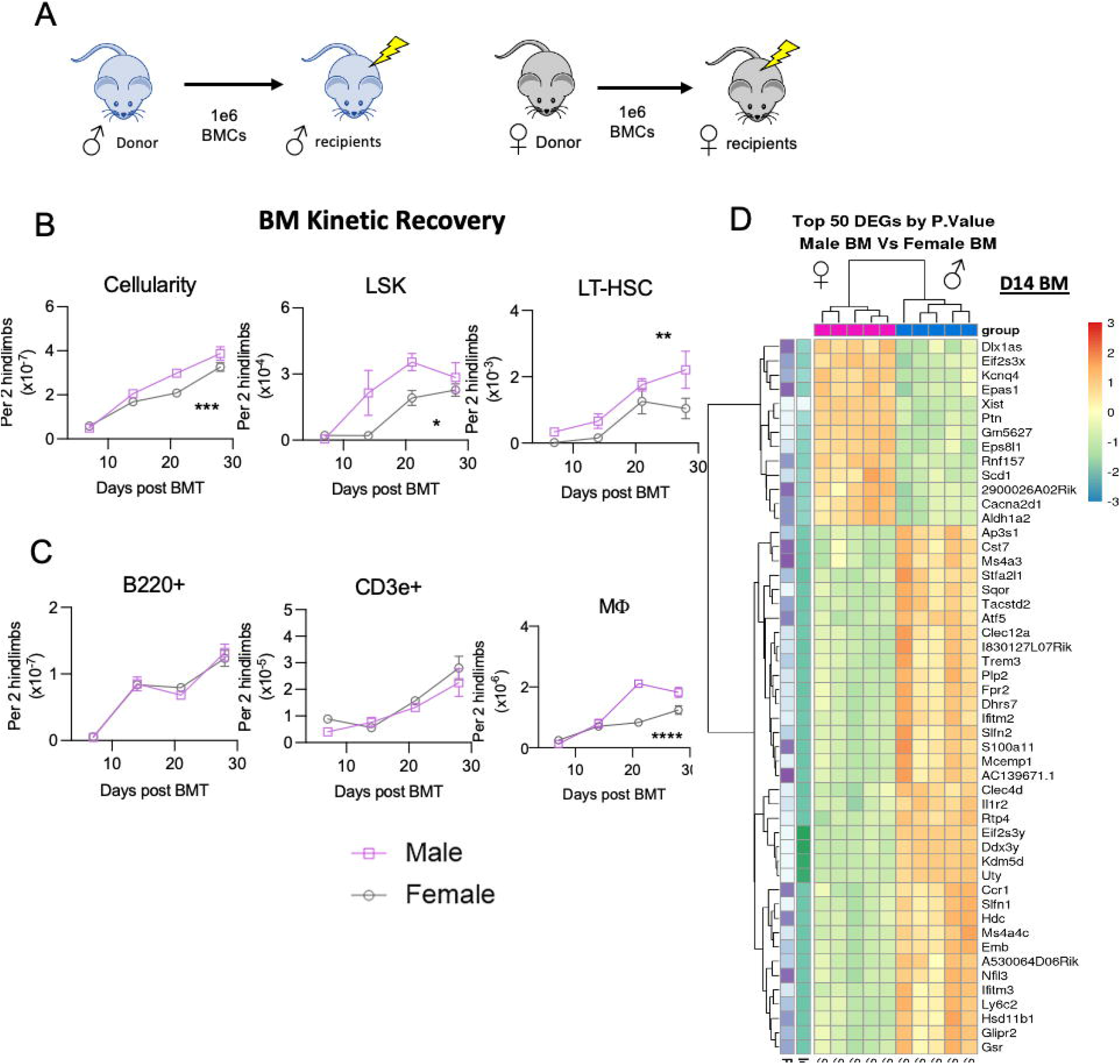
Male mice display accelerated medullary hematopoietic recovery following HST-. A. Schematic depicting transplant conditions. B. BM cellularity, and quantification of lineage-cKit+Sca1+ (LSK) and LSK CD150+CD48- (LT-HSC) was obtained on cohorts of mice sacrificed on Days 7, 14, 21, and 28 post transplant. N=5 mice/arm for D7 and D28, N=10 mice/arm for D14, D21. C. Quantification of BM B-cells (B220+), T-cells (CD3e+), and Macrophages (F4/80+) on cohorts described in B. N=5 mice/arm for D7 and D28, N=10 mice/arm for D14, D21. N=-5 mice/arm for Macrophages. Error bars represent SEM. 2-way ANOVA was performed and asterisks denote male vs. female. D. Heat map generated from bulk RNA sequencing on total BM populations of D14 post HST mice. Top 50 differentially expressed genes ranked by p-value. N=5 mice/arm.

The splenic compartment also showed robust recovery advantages in the M➔M cohort compared to F➔F as evidenced by significant increases in total cellularity at D7 (37.6M/mouse vs. 12.7M), D14 (47M/mouse vs. 29.3M), D21 (79.5M/mouse vs. 50.4M), and D28(70.9M/mouse vs. 46.2M). We also observed significant increases in the LSK population at D7 (132K vs. 45K), D14 (60.6K vs. 15K), and D21(100K vs. 32.6K) (**Figure 5B**). Recovery of the B, T, and macrophage populations in the spleens of M➔M vs. F➔F mice was dramatically different, with M➔M mice displaying higher numbers of B-cells particularly at D21 (5.2M/spleen vs. 3M) and D28 (3.9M/spleen vs. 2.1M), and macrophages at each timepoint, particularly the D21 (3.8M/spleen vs. 1.8M) and D28 ( 3.2M/spleen vs. 1.7M) (**Figure 5C**). RNA sequencing results also demonstrate splenic sex differences between genes associated with hematopoietic regeneration including downregulation of *Cfh* (potentially reducing splenic inflammation and autoimmunity ^29^*), Ptprd (*inhibition of Ptp has been demonstrated to promote regeneration ^30^*), Cxcl12 (*suggesting enhanced HSC mobilization*), Epas1(also* observed in the BM, which may promote regeneration*), Lepr (which marks long-term HSCs ^31^*), and *Malat1* (which is downregulated upon hematopoietic differentiation ^32^) in male compared to female recipients (**Figure 5D).**

**Figure 5:**
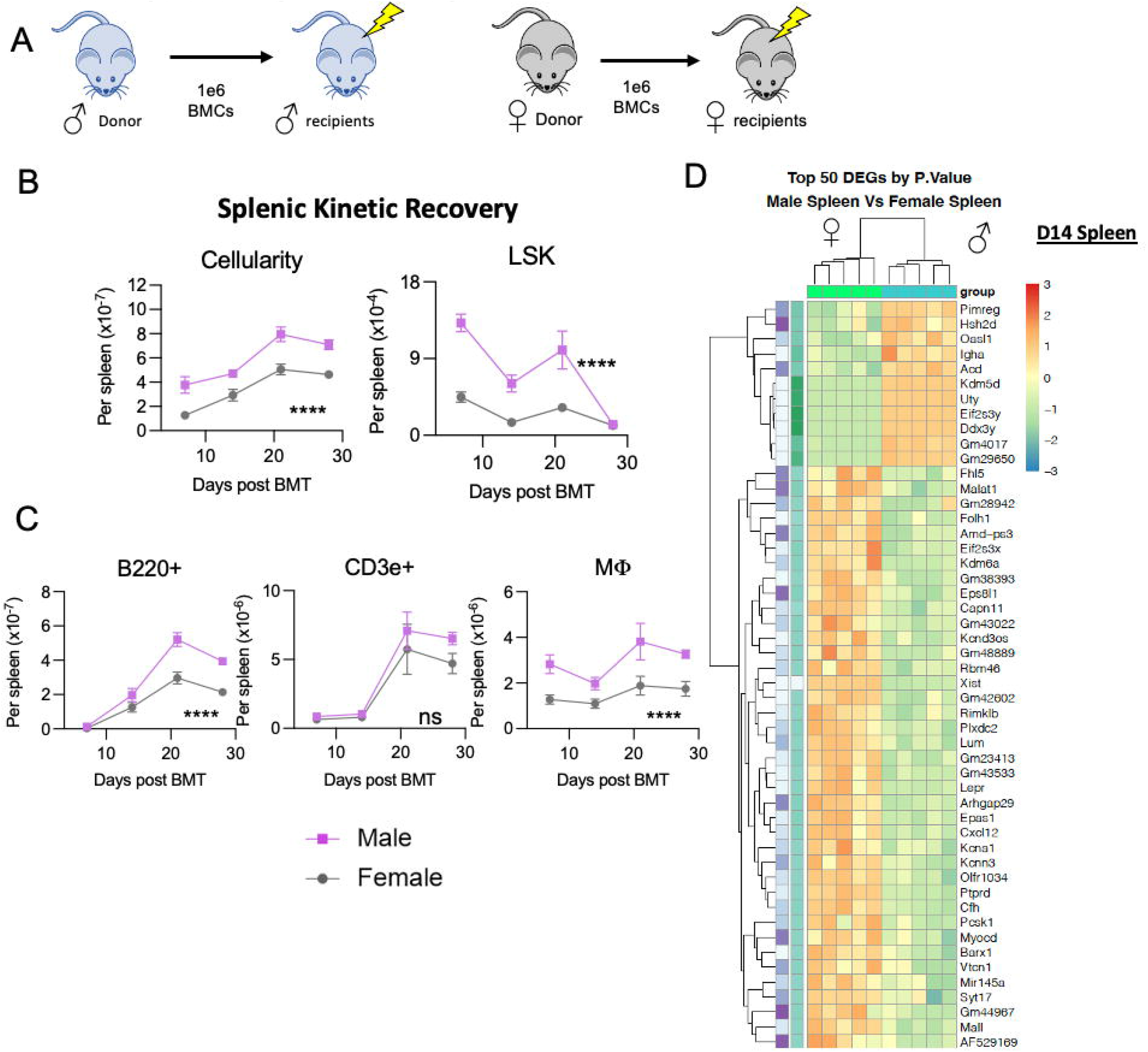
Male mice exhibit robust splenic extramedullary hematopoiesis post-transplant-. A. Schematic depicting transplant conditions. B. Splenic cellularity, and quantification of lineage-cKit+Sca1+ (LSK) and LSK CD150+CD48- (LT-HSC) was obtained on cohorts of mice sacrificed on Days 7, 14, 21, and 28 post transplant. C. Quantification of splenic B-cells (B220+), T-cells (CD3e+), and Macrophages (F4/80+) on cohorts described in B. N=5 mice/arm for D7 and D28, N=10 mice/arm for D14, D21. N=-5 mice/arm for Macrophages. Error bars represent SEM. 2-way ANOVA was performed and asterisks denote male vs. female. D. Heat map generated from bulk RNA sequencing on total BM populations of D14 post HST mice. Top 50 differentially expressed genes ranked by p-value. N=5 mice/arm.

### The male microenvironment drives hematopoietic recovery independent of HSC donor sex

To distinguish the roles of donor HSCs vs. the recipient microenvironment in contributing to the sexual dimorphism of HST regeneration, we established mismatch transplants with each combination of donor versus recipient (M➔M, M->F, F->M, F➔F). We sacrificed these cohorts at D14 post HST, as this marked a time point when HSCs and HSPCs exhibited sex-dependent recovery differences that preceded effects on mature cell types, and characterized the HSPC population in the BM and splenic microenvironments. Notably, we observed that F->M mice, which consisted of female donor BM transplanted into a male recipient, recovered at an accelerated pace compared to M->F mice, which consisted of male donor BM transplanted into a female recipient, and that the male microenvironment was generally sufficient to recapitulate the enhanced recovery observed in the M➔M scenario. This included significant increases in BM cellularity (20.3M/mouse vs. 17.4M) and trends toward increased BM LSK (19.5K/mouse vs. 11.4K/mouse) and LT-HSC (288/mouse vs. 100) numbers). In the spleen we observed that the F➔M group also displayed accelerated recovery compared to M➔F mice as shown by increased D14 splenic cellularity (34.3M/mouse vs. 25.3M) and LSK (67.3K/mouse vs. 26.5K) numbers (**Figure 6).** This data suggests that the male microenvironment is responsible for the accelerated recovery post HST, and that this phenotype is independent of donor cell sex.

**Figure 6:**
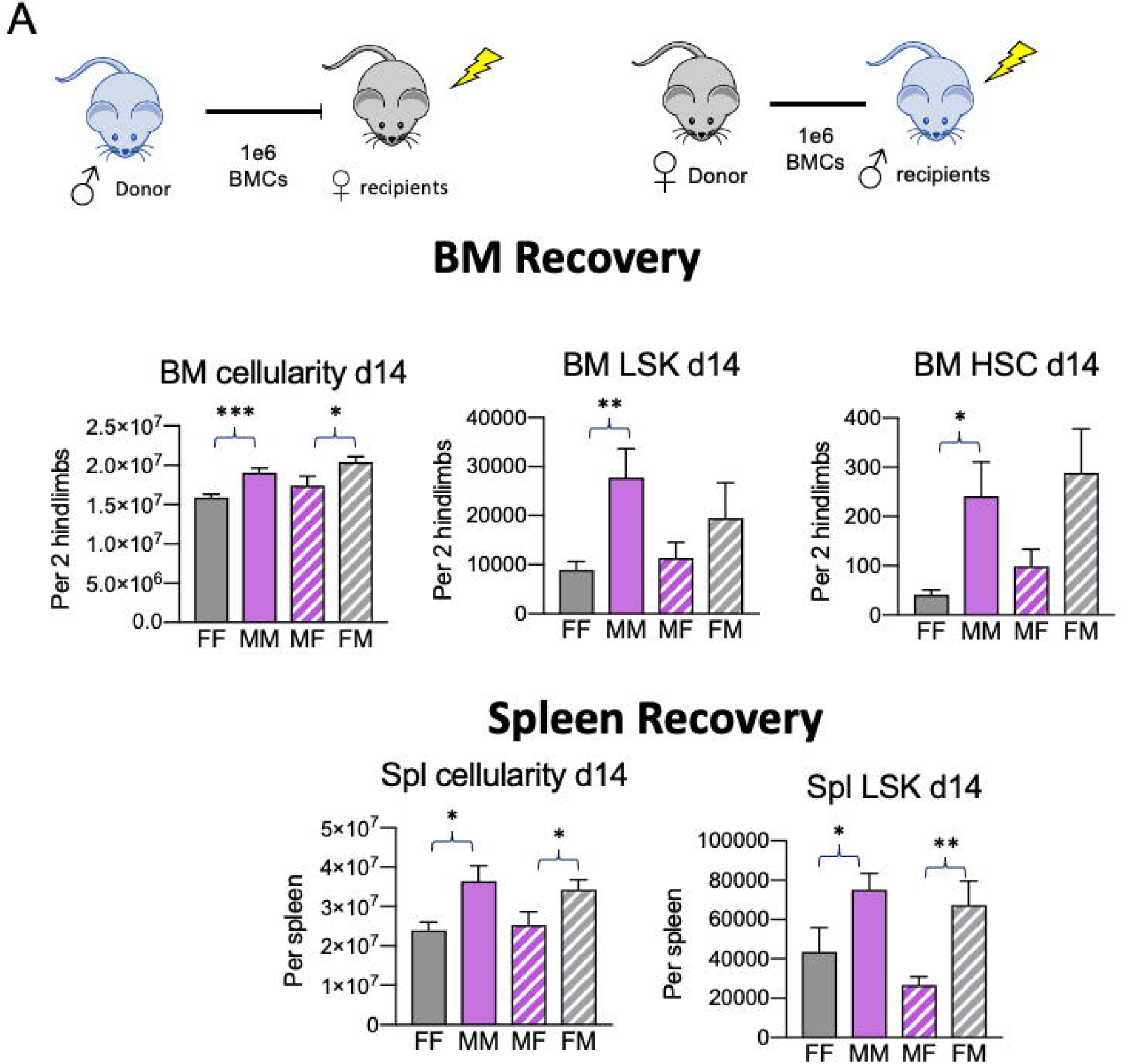
The male microenvironment drives hematopoietic recovery independent of donor HSC sex-. A. Schematic depicting transplant conditions. 1e6 BM cells from male donors were transplanted into lethally irradiated (10Gy) female recipients, and the equivalent performed using female donors and male recipients. B. BM and splenic cellularity, and quantification of lineage-cKit+Sca1+ (LSK) and LSK CD150+CD48- (LT-HSC) was obtained from mice sacrificed on Day 14 post transplant. N=10 mice/arm. Error bars represent SEM. \*\**P* < 0.01. Student’s t-test used for all analyses.

### Estradiol supplementation to BM cultures significantly increases long-term HSCs

We next sought to elucidate a potential role for sex hormones in mediating our observed differences in steady state composition and post-transplant regeneration rates between male and female mice. Coviello *et. al* have previously demonstrated that male patients receiving weekly doses of testosterone (T) display a dose dependent stimulation of erythropoiesis and that this stimulation is more pronounced in elderly males^33 34^. This data suggests that testosterone yields a lineage specific hematopoietic burst that results in an expansion of erythroid populations. Multiple groups have also demonstrated a role for 17β estradiol (E2) in HSC regulation. Illing *et. al* published that E2 administration to mice resulted in increased HSC numbers in the vascular (but not endosteal) hematopoietic niche by increasing HSC cell cycle entry ^35^, and Nakada *et. al* subsequently showed that elevated E2 levels during pregnancy resulted in increased HSC division, HSC frequency, cellularity, and erythropoiesis in the spleen ^4^. Here we established cultures of total BM supplemented with increasing doses of either E2 or T (3pg/mL, 30pg/mL, 300pg/mL) and harvested these cultures at D5 following treatment.

Assessment of the LSK and LT-HSC (SLAM) populations demonstrate that E2 supplementation to both male and female BM results in a significant decrease in the frequency of LSK cells. 30pg/mL E2 resulted in ~2-fold decrease in LSK while 300pg/mL led to a ~3-fold decline. Surprisingly, this was coupled with significant increases in the frequency of SLAM cells in culture which suggests that sex steroids may alter the *ex vivo* proliferation and differentiation capacity of HSCs (**Figure 7**). We also demonstrated that this effect was specific to E2 treatment, as T supplementation to BM cultures resulted in no significant alterations to either the LSK or SLAM compartments (**Supplemental Figure 2**).

**Figure 7:**
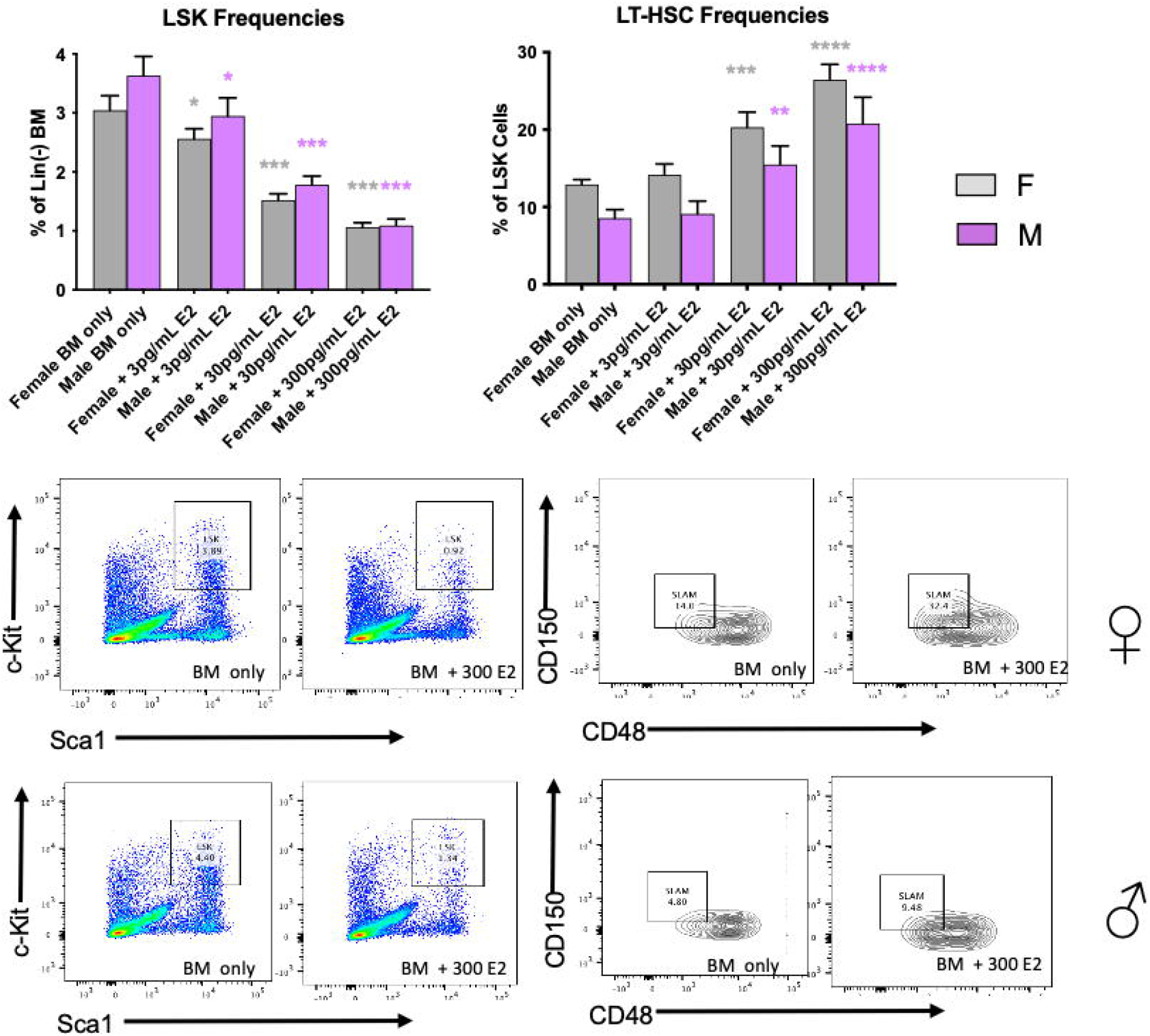
Estradiol supplementation to BM cultures significantly increases long-term HSCs-. A. Frequencies of lineage-cKit+Sca1+ (LSK) and LSK CD150+CD48- (LT-HSC) following 5 days of total male and female BM cultured with the indicated doses of Estradiol (E2). B. Representative flow cytometric plots depicting BM LSK and LT-HSC populations on Day 5 cultures of BM only versus BM + 300 pg/mL E2. n=7 mice.arm. Error bars represent SEM. \*\**P* < 0.01. Student’s t-test used for all analyses and represent each experimental arm again untreated BM.

## Discussion

Recent work studying sex-differences in hematopoiesis has elucidated an important role for sex in mediating (1) obesity induced inflammatory response-with males demonstrating enhanced myelopoiesis and inflammation during obesity ^36^, (2) HSC self-renewal and proliferation capacity-with estrogen promoting HSC division, self-renewal capacity, and erythroid output ^4^, (3) aging hematopoiesis- with male HSCs exhibiting an earlier phenotypic and functional decline than females ^37^, (4) incidence of infectious and autoimmune diseases - with more robust interferon responses and altered macrophage gene expression in females proposed as being causative^38^. Further understanding the role of sex in additional aspects of hematopoiesis is warranted-particularly as sexual dimorphism has been demonstrated as an important player in pathogenesis and treatment options for a number of diseases including many cancers ^39 40^, diabetes ^41^, cardiovascular disease ^42^, and others.

In this manuscript we begin to both phenotypically and mechanistically characterize sexual dimorphism associated with steady state and regenerative hematopoiesis. In particular we identified (1) that male mice at steady state contain an increased number of LSK cells and trend towards reduced LT-HSCs, (2) that steady state male and female mice uniquely express factors critical to the hematopoietic niche, with males demonstrating a BM signature consistent with increased HSC production and proliferation, (3) that these gene expression signatures correlate with accelerated medullary and extra-medullary recovery upon syngeneic transplantation of male donor cells into male recipients compared to female donor cells into female recipients, (4) that the male microenvironment is responsible for this observed accelerated recovery, as female donor BM transplanted into male recipients recovered post HST faster than male BM transplanted into female recipients, and (5) that E2 supplementation to BM cultures reduces the frequency of LSK cells but significantly increases frequencies of long-term SLAM cells.

We thus hypothesize that sex-specific cellular and molecular signaling occurs during hematopoietic regeneration. We also implicate steroid sex hormones in determining regeneration rates and propose that E2 promotes long-term HSC populations at the expense of the more differentiated short-term LSK cells. This may help explain our data from **Figures 4 and 5,** if we hypothesize that increased E2 production in females during HST limits LT-HSC proliferation and progenitor differentiation, thus restricting early hematologic recovery in female versus male recipients. On the other hand, we expect that this effect of E2 on LT-HSCs will provide long-term advantages over the lifetime of the recipient. This hypothesis is intriguing and requires further study.

This work has also uncovered additional questions that require further study, including:

What are the unique sex-specific roles of the donor HSCs versus recipient stromal cells on hematopoietic engraftment, proliferation, and long term maintenance following HST?
What are the sex-dependent genetic and phenotypic differences in hematopoietic regeneration following myeloablation?
What roles do steroid sex hormones play on transplantation outcomes?
Whether steroid sex hormone differences underlie the differential expression of HSC niche genes in the BM and spleen.

Elucidating additional sex-specific mechanisms of regenerative hematopoiesis has implications for the approximately ~25,000 HST are performed each year in the United States to treat conditions including plasma cell dyscarsias, non-Hodgkin’s lymphoma, acute myeloid leukemia, myelodysplastic syndromes, and acute lymphocytic leukemia. While HST is curative in many settings and capable of significantly extending survival in others, the severe complications associated with the procedure have limited its utility and narrowed the potential patient pool^43,44^. By deepening our understanding of factors regulating hematopoiesis and hematopoietic stem cell function, we have enormous potential to expand the utility of HST.

## Acknowledgments

This work was supported by NIH grants R00 HL135740, R01 HL158801-01, T32 EB005583, and by the Radiation Resources Core Facility, the Hematopoietic Biorepository and Cellular Therapy Core Facility, the Applied Functional Genomics Core, and the Cytometry & Imaging Microscopy Core Facility of the Case Comprehensive Cancer Center (P30CA043703). This work was also supported by a generous award from the Ohio Cancer Research Foundation.

## Author Contributions

Julianne N.P. Smith: Conception and design, Data collection/interpretation, Manuscript editing Brittany A. Cordova: Data collection/interpretation

Brian Richardson: Data interpretation

Kelsey F. Christo: Data collection

Jordan Campanelli: Data collection/interpretation

Alyssia V. Broncano: Data collection

Jonathan Chen: Data collection

Juyeun Lee: Data interpretation, Manuscript editing

Scott J. Cameron: Data interpretation, Manuscript editing

Justin D. Lathia: Data interpretation, Manuscript editing

Wendy A. Goodman: Data collection/interpretation, Manuscript editing

Mark J. Cameron: Data interpretation, Manuscript editing

Amar B. Desai: Conception and design, Data collection/interpretation, manuscript writing, Final approval of manuscript

