## Supplemental Data for "Sex-specific niche signaling contributes to sexual dimorphism following stem cell transplantation"

### Slide 1
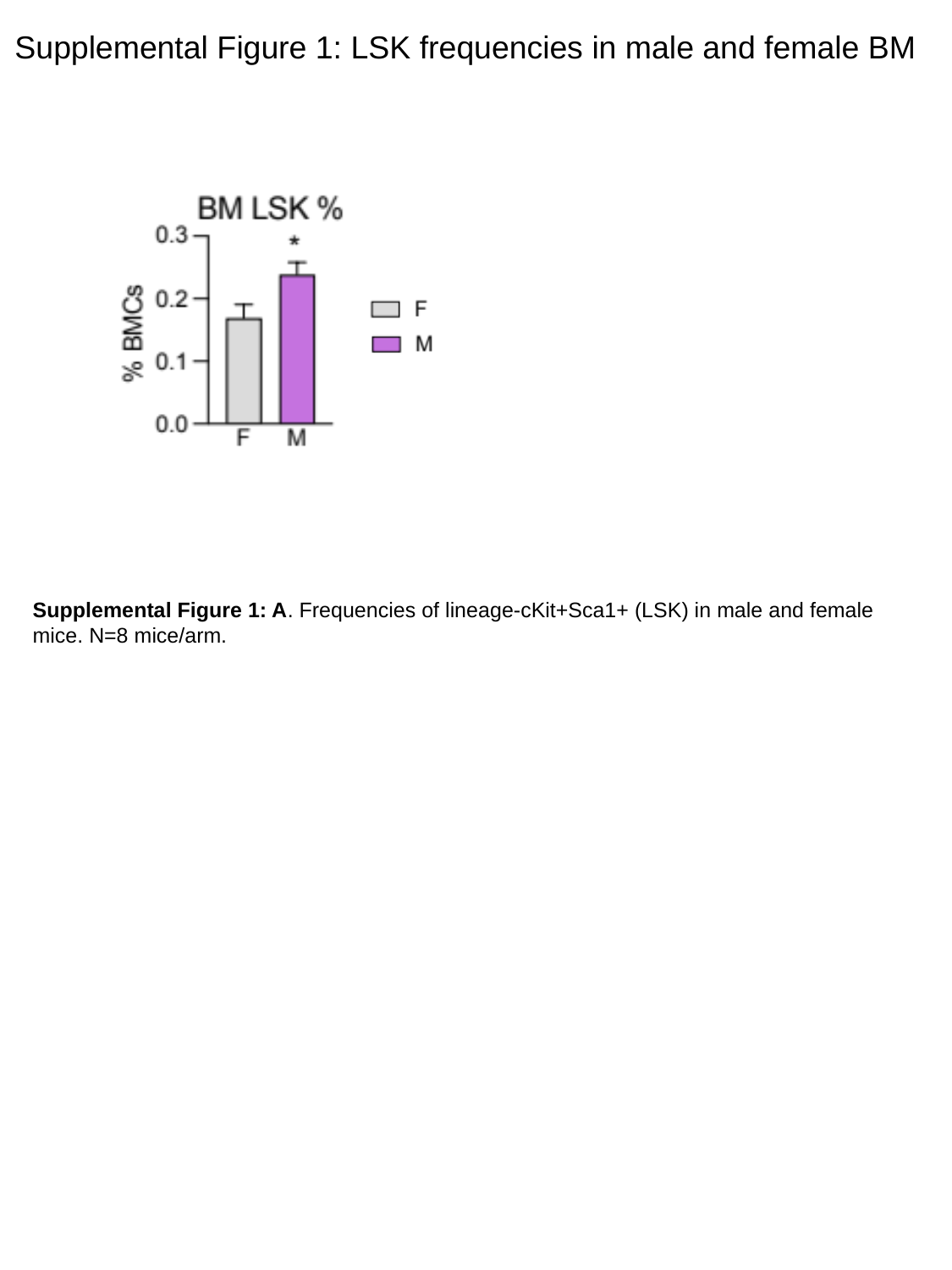

Supplemental Figure 1: LSK frequencies in male and female BM
Supplemental Figure 1: A. Frequencies of lineage-cKit+Sca1+ (LSK) in male and female mice. N=8 mice/arm.

### Slide 2
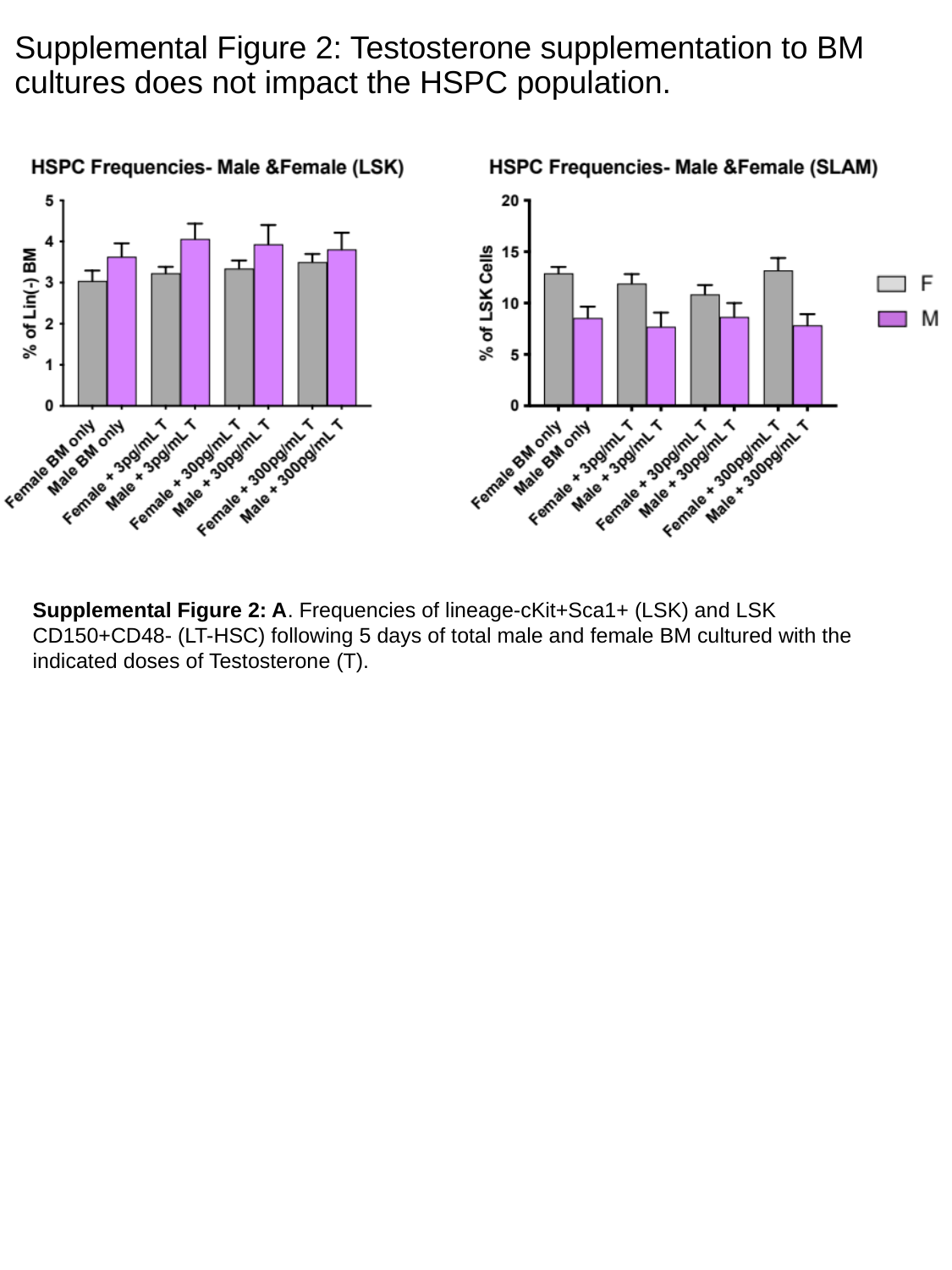

Supplemental Figure 2: Testosterone supplementation to BM cultures does not impact the HSPC population.
Supplemental Figure 2: A. Frequencies of lineage-cKit+Sca1+ (LSK) and LSK CD150+CD48- (LT-HSC) following 5 days of total male and female BM cultured with the indicated doses of Testosterone (T).
